# Development of a mutant aerosolized ACE2 that neutralizes SARS-CoV-2 in vivo

**DOI:** 10.1101/2023.09.26.559550

**Authors:** Daniel L. Kober, Marley C. Caballero Van Dyke, Jennifer L. Eitson, Ian N. Boys, Matthew B. McDougal, Daniel M. Rosenbaum, John W. Schoggins

## Abstract

The rapid evolution of SARS-CoV-2 variants highlights the need for new therapies to prevent disease spread. SARS-CoV-2, like SARS-CoV-1, uses the human cell surface protein angiotensin-converting enzyme 2 (ACE2) as its native receptor. Here, we design and characterize a mutant ACE2 that enables rapid affinity purification of a dimeric protein by altering the active site to prevent autoproteolytic digestion of a C-terminal His_10_ epitope tag. In cultured cells, mutant ACE2 competitively inhibits lentiviral vectors pseudotyped with spike from multiple SARS-CoV-2 variants, and infectious SARS-CoV-2. Moreover, the protein can be nebulized and retains virus-binding properties. We developed a system for delivery of aerosolized ACE2 to K18-hACE2 mice and demonstrate protection by our modified ACE2 when delivered as a prophylactic agent. These results show proof-of-concept for an aerosolized delivery method to evaluate anti-SARS-CoV-2 agents *in vivo* and suggest a new tool in the ongoing fight against SARS-CoV-2 and other ACE2-dependent viruses.

## Introduction

The severe acute respiratory syndrome coronavirus 2 (SARS-CoV-2) causes coronavirus-induced disease 19 (COVID-19) (Zhu et al. 2020). The rapid, ongoing evolution of SARS-CoV-2 has produced variants of concern, such as omicron, which can escape immune responses and evade antibody therapies (Hoffmann et al. 2022; Liu et al. 2022; Cele et al. 2022). A pan-coronavirus therapy that protects against existing and emerging coronaviruses would be a valuable tool with immediate and long-term potential utility, however this goal is challenging since such therapeutics would have to neutralize as-yet-unevolved virions. The cell-surface receptor for SARS-CoV-2 is human angiotensin-converting enzyme 2 (ACE2) (Hoffmann et al. 2020; Yan et al. 2020), a receptor that has been exploited by other pathogenic coronaviruses including SARS-CoV-1 (Li et al. 2003). ACE2 is a zinc carboxypeptidase (Tipnis et al. 2000; Donoghue et al. 2000) that converts the 8-residue peptide Angiotensin II (‘Ang II’) to An-(1-7). Ang II is produced by angiotensin-converting enzyme (ACE), and the opposing activities of ACE and ACE2 regulate the renin-angiotensin system (RAS) (Turner 2015). Delivery of an enzymatically inactive ACE2 as an anti-SARS-CoV-2 agent would potentially be less disruptive to this regulatory system. Soluble derivatives of ACE2 have been proposed for treating and preventing SARS-CoV-2, and are being explored in pre-clinical studies (Shoemaker et al. 2022; Haschke et al. 2013; Monteil et al. 2021) and clinical settings (Zoufaly et al. 2020). The ideal formulation of such an ACE2 would enable rapid, large-scale purification of a protein that retains anti-viral efficacy following nebulization for delivery into the respiratory system. To our knowledge, nebulized delivery of a soluble ACE2 that prevents SARS-CoV-2 infection in animals has not been demonstrated.

Cryo-EM studies of full-length ACE2 show the protein is a dimer on the cell surface, where it binds the SARS-CoV-2 receptor binding domain (RBD) (Fig 1A). The ectodomain of ACE2, corresponding to residues 19-740, can be subdivided into two domains, the Enzyme domain and the Neck domain. Most of the dimer contacts are between the two Neck domains of the ACE2 monomers, with additional interactions occurring between the ACE2 Enzyme domains. Each spike RBD engages with an ACE2 monomer away from the dimer interface. Similarly, the ACE2 active site is not part of the dimer interface or the RBD-binding surface (Yan et al. 2020) (Fig 1A), and disrupting the ACE2 active site would not be expected to disrupt SARS2-CoV-2 binding to ACE2.

**Figure 1.**
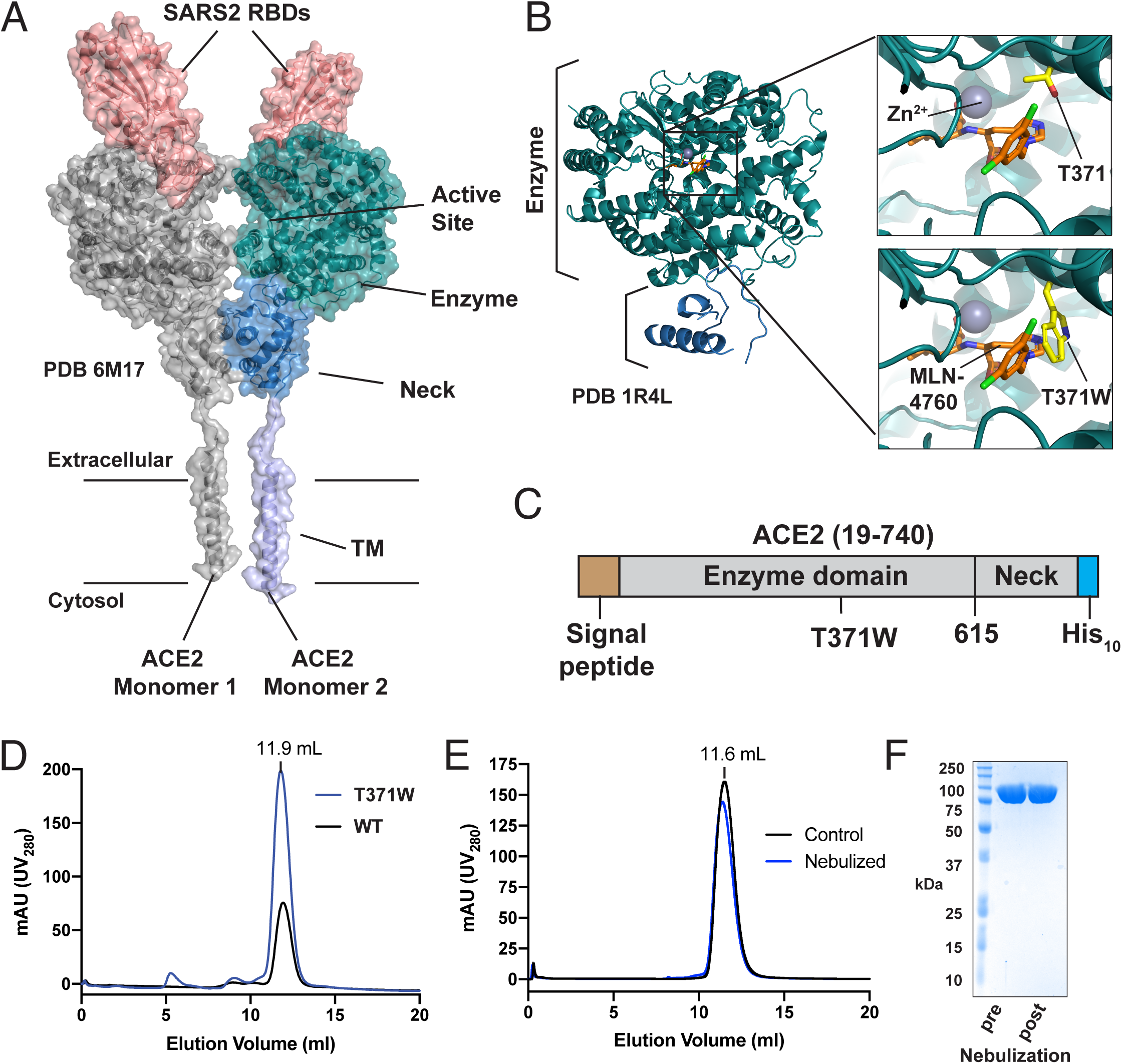
Design and characterization of a mutant dimeric ACE2. A) Cryo-EM structure of full-length ACE2 bound to SARS2 RBDs (PDB 6M17). One ACE2 monomer is depicted in grey. The other ACE2 monomer is depicted in purple, blue, and teal for the transmembrane (TM), Neck, and Enzyme domains, respectively. Bound RBDs are shown in red. Approximate position of the membrane boundaries are indicated. B) Structure-guided design of the active site T371W mutant. The crystal structure of ACE2 ectodomain bound to MLN-4760 (PDB 1R4L) is shown on the left, with the domains colored as in (A). Insets show active site with MLN-4760, T371, and T371W mutations depicted with sticks. Zn^2+^ metal shown with grey sphere. C) Schematic for the soluble ACE2 ectodomain construct for insect cell expression D) Gel filtration traces for ACE2 WT and ACE2^T371W^ ectodomains grown in culture supplemented with 1 mM EDTA. NiNTA-purified protein samples were subjected to gel filtration over a superdex 200 column. E) Gel filtration traces for ACE2^T371W^ ectodomain samples treated with 1% w/v PEG-8000 collected before nebulization (Control) or after nebulization (Nebulized). F) Coomassie-stained SDS-PAGE analysis of the samples from (E).

Here, we developed a version of ACE2 with a space-filling mutation to the catalytic pocket that greatly reduces the protease activity of ACE2 and facilitates rapid purification of the secreted protein using one-step affinity chromatography. This protein retains the ability to bind the RBD of SARS-CoV-2 and SARS-CoV-1 and prevents viral infection *in vitro*. Aerosolized ACE2 delivered using a nebulizer apparatus prevented disease in K18-hACE2 mice. These data demonstrate proof-of-concept for the use of a modified, catalytically impaired ACE2 to prevent COVID-19 through aerosol delivery.

## Results

To explore the use of nebulized ACE2 as a candidate for anti-COVID-19 therapy, we designed a version of soluble ACE2 that retained the virus-neutralizing properties of endogenous soluble ACE2 and was engineered to enable simple and rapid purification. Using the crystal structure of the ACE2 ectodomain bound to the inhibitor MLN-4760 as a guide (Towler et al. 2004), we designed a space-filling mutation within the active site, T371W, which is predicted to clash with MLN-4760 (Fig 1B). This residue is not catalytic and does not participate in Zn^2+^ coordination. We reasoned that this mutation would sterically block substrate binding, which could prove advantageous for purification and eliminate enzymatic activity for the therapeutic candidate.

We generated baculovirus encoding a melittin signal peptide, the complete human ACE2 ectodomain (residues 19-740), and a C-terminal His_10_ tag (Fig 1C). Comparing expression of ACE2 WT and ACE2^T371W^ in Sf9 insect cells using anti-His and anti-ACE2 antibodies, we observed similar levels of secreted soluble ACE2 in the Sf9-conditioned supernatant (via anti-ACE2 antibody), however the ACE2 WT culture had greatly decreased signal for the C-terminal His_10_ tag compared to ACE2^T371W^. Because ACE2 is a zinc carboxy-peptidase, we reasoned that the soluble ACE2 ectodomains may proteolyze their own C-terminal His_10_ tag. Indeed, supplementing the Sf9 culture media with 1 mM EDTA rescued levels of the ACE2 His_10_ tag in the culture media (Appendix Fig S1A). Previous structural biology studies on ACE2 with purification by nickel-nitrilotriacetic acid (NiNTA) resin used C-terminally tagged ACE2 consisting of the enzyme domain (residues 19-615) without the neck domain (residues 616-740) (Kirchdoerfer et al. 2018; Li 2008; Li et al. 2005; Song et al. 2018; Wu et al. 2012; Lan et al. 2020). The only structure containing the full soluble ectodomain utilized protein purified by a multi-step tag-less biochemical protocol (Towler et al. 2004). While the Neck domain was not fully modeled in this study due to poor electron density, symmetry expansion of PDB 1R4L shows the same dimeric interface between Necks characterized in the transmembrane ACE2 structure using cryo-EM (Yan et al. 2020). We speculate that the extended Neck domain may allow cleavage of the C-terminal His tag by ACE2, whether in trans or in cis. Regardless, preserving the ACE2 Neck domain is likely critical to preserve the dimeric form observed in full-length ACE2 (Yan et al. 2020).

We hypothesized that our T371W mutation might permit rapid and efficient large-scale purification of His-tagged, dimeric ACE2 ectodomain. To obtain comparable ACE2 WT and ACE2^T371W^ samples (avoiding C-terminal proteolysis), we supplemented the Sf9 culture with 1 mM EDTA and conducted His-tag purification of secreted protein using an EDTA-resistant IMAC resin. Both ACE2 WT and ACE2^T371W^ ectodomains could be purified in this manner, albeit with yields of ≤ 1 milligram per liter of culture. These proteins showed similar behavior on gel filtration chromatography (Fig 1D). We characterized the enzymatic activity of these proteins using an established ACE2 fluorescent self-quenched peptide cleavage assay (Vickers et al. 2002) where the substrate peptide Mca-APK(Dnp) is incubated with the enzyme and Mca fluorescence is monitored. Whereas ACE2 WT showed activity when supplemented with 10 µM ZnCl_2_, ACE2^T371W^ remained largely inactive (Appendix Fig S1B). Moreover, WT ACE2 was inhibited with 10 µM MLN-4760, but the residual activity of ACE2^T371W^ was not further inhibited by MLN-4760 (Appendix Fig S1C). These measurements confirm that the T371W mutation disrupts the ACE2 active site.

Next, we characterized the solution behavior of ACE2^T371W^. For all the remaining experiments, ACE2^T371W^ was expressed without supplementing the culture with EDTA, as this was no longer needed to protect the C-terminal His_10_ tag from proteolysis. In this format, we routinely obtained greatly enhanced yields of 10 milligrams per liter culture without further optimization. As expected, ACE2^T371W^ contains N-linked glycosylations (Appendix Fig S2A). The purified protein is dimeric as judged by size exclusion chromatography-multi angle light scattering (SEC-MALS) analysis (predicted dimer MW = 172 kDa, experimental MW = 163 kDa) (Appendix Fig S2B). The protein shows good thermal stability by differential scanning fluorimetry (DSF) with a T_m_ of ∼48°C. ACE2^T371W^ is marginally less stable when the N-linked glycans are removed using PNGaseF (Appendix Fig S2C-D). We also measured the denaturation of the protein using circular dichroism spectroscopy (CD), which showed a single transition at ∼50°C. Interestingly, the protein did not lose all secondary structure, even with temperatures up to 95°C (Appendix Fig S2E).

Next, we tested whether the dimeric ACE2^T371W^ survives nebulization. ACE2^T371W^ collected before or after nebulization with 1% w/v PEG-8000 as excipient (hereafter soluble [sACE2^T371W^] and nebulized [nebACE2 ^T371W^], respectively) displayed identical gel filtration profiles, and nebACE2^T371W^ remained intact as shown by Coomassie-stained SDS-PAGE (Fig 1E-F). To test whether nebulized ACE2^T371W^ retains binding to the RBDs of SARS-CoV-1 and SARS-CoV-2 spike proteins, we employed isothermal titration calorimetry (ITC). The RBDs of these viruses were expressed as secreted FLAG-tagged proteins from Sf9 insect cells and purified by anti-FLAG chromatography and gel filtration (Fig 2A). ITC experiments were conducted to measure the heat of binding between ACE2^T371W^ and the RBDs (Fig 2B). Strong binding isotherms were observed for binding to both SARS-CoV and SARS-CoV2 RBDs, and the affinity constants for sACE2^T371W^ and nebACE2^T371W^ were nearly identical (Fig 2C). Notably, these values are consistent with the K_D_=15 nM reported using surface plasmon resonance (SPR) measurements on SARS2 and ACE2 ectodomains (Wrapp et al. 2020).

**Figure 2.**
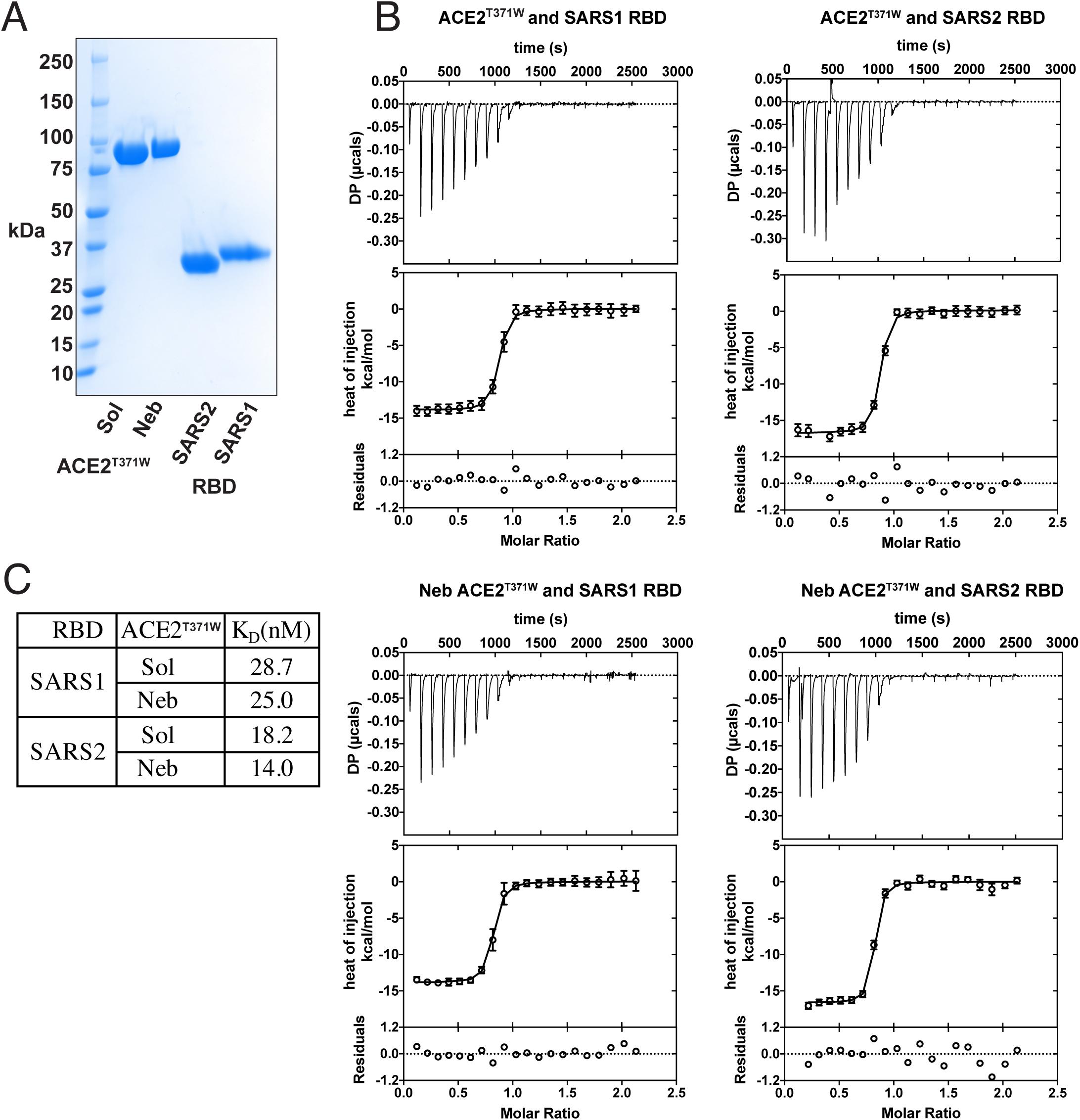
Nebulized ACE2^T371W^ retains binding to SARS1 and SARS2 RBDs. A) Coomassie-stained SDS-PAGE analysis of protein samples for ITC experiments. B) Isotherms for ACE2^T371W^ and RBD isothermal titration calorimetry experiments. upper panel shows the SVD-corrected thermograms from NITPIC; the middle panel displays the integrated data points with their respective estimated error bars, and the solid line results from the fit. The lower panel depicts the residuals between the data and the fit lines. C) Affinity values derived from ITC data. The binding reactions between sACE2 and the SARS2 RBD or SARS1 RBD were measured from four or two independent purifications, respectively. The binding reactions between nebACE2 and the SARS1 and SARS2 RBDs were measured from one purification.

We next tested whether ACE2^T371W^ inhibits viral infection, starting with a tractable assay that could be performed in Biosafety Level 2 conditions. We used replication-defective HIV-1-based lentivirus particles pseudotyped with the spike protein from several SARS-CoV-2 variants (G614, alpha, beta, gamma, delta, omicron), or with the glycoprotein from vesicular stomatitis virus (VSV-G) as a control. The lentiviruses encode firefly luciferase as a readout of viral transduction. Huh7.5 cells ectopically expressing ACE2 were transduced with lentiviruses in the presence of sACE2^T371W^. The next day, luciferase signals from cell lysates were quantified, and data were normalized to the levels of luciferase generated from untreated, transduced cells. In the presence of sACE2^T371W^ concentrations ranging from 333nM to 0.15nM, VSV-G-pseudotyped lentivirus was not inhibited (Fig 3A). In contrast, spike-pseudotyped lentiviruses incubated with sACE2^T371W^ were inhibited in a dose-dependent manner. Inhibitory effects varied across the spike variants, with suppression ranging from 65%-95% at the maximum dose of 333nM sACE2^T371W^.

**Figure 3.**
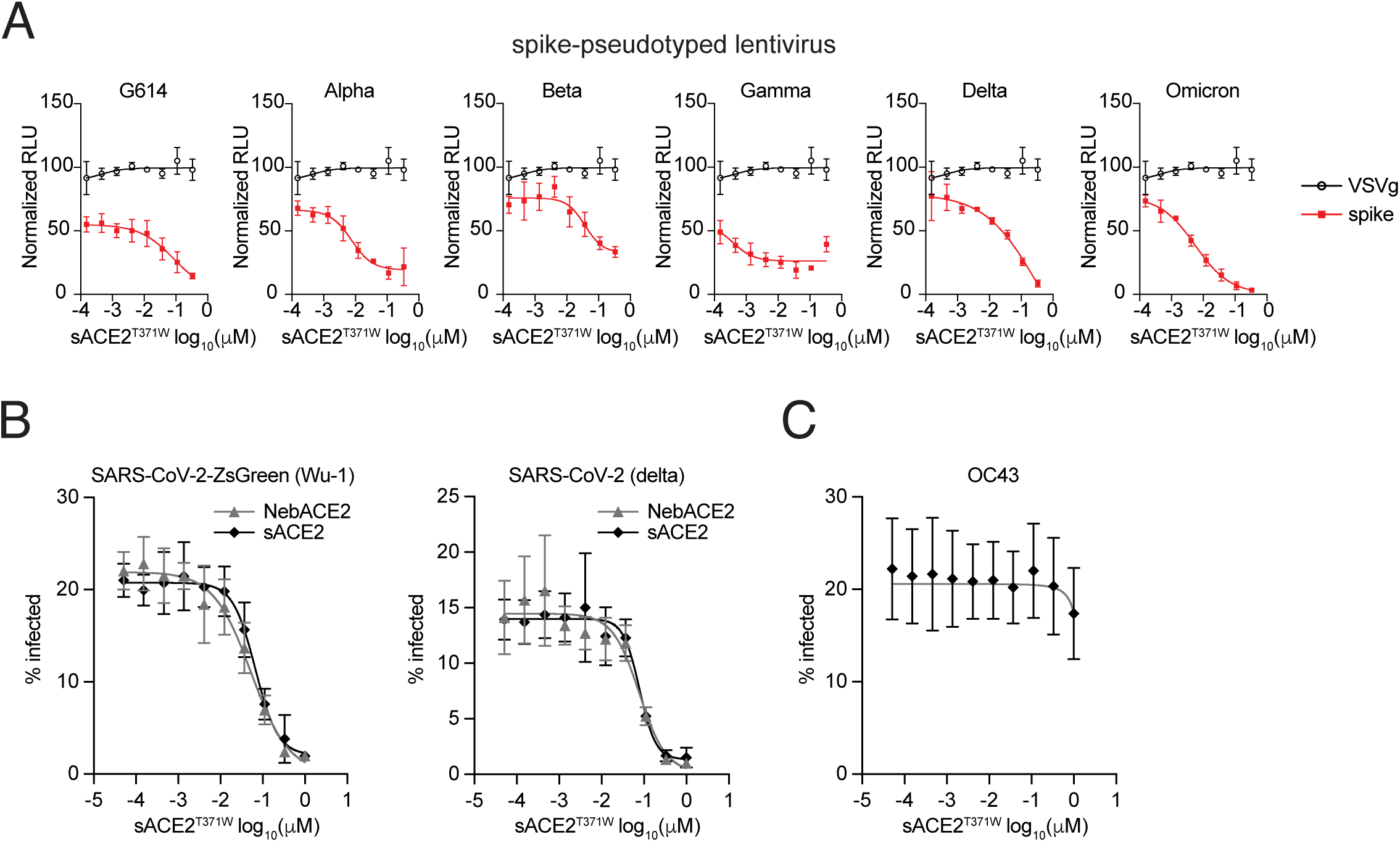
sACE2^T371W^ and nebACE2^T371W^ inhibit spike-pseudotyped lentivirus and SARS-CoV-2. A) Infectivity of luciferase-expressing SARS-CoV-2 spike-psuedotyped lentiviruses (red) or VSV-G-psuedotyped lentivirus (black) in the presence of varying concentrations of sACE2^T371W^. Transduction assays were performed in Huh7.5-mutACE2 cells. N= 3 biological replicates. B) Infectivity of SARS-CoV-2-Wu-1-zsGreen or delta variant in the presence of varying concentrations of sACE2^T371W^ (black) or nebACE2^T371W^ (gray). Infection assays were performed in A549-TMPRSS2-ACE2 and viral infectivity was quantified by flow cytometry. N= 3 biological replicates. C) Infectivity of human coronavirus OC43 in the presence of varying concentrations of sACE2^T371W^. Infection assays were performed in A549-TMPRSS2-ACE2 and viral infectivity was quantified by flow cytometry. N= 3 biological replicates.

We next tested the inhibitory potential of ACE2^T371W^ against authentic SARS-CoV-2 in cultured cells in a Biosafety Level 3 setting. Over a 10-point dilution series starting at 1μM, we found that sACE2^T371W^ strongly inhibited a ZsGreen-expressing recombinant SARS-CoV-2 (Wu-1 strain) and a clinical isolate of the delta variant (Fig 3B). The human coronavirus OC43, which uses sialic acid instead of ACE2 for entry (Vlasak et al. 1988), was not inhibited by sACE2^T371W^ (Fig 3C). Notably, SARS-CoV-2 was also inhibited by the nebulized form of ACE2^T371W^ (Fig 3B). This prompted us to examine whether aerosolized ACE2^T371W^ could be administered to mice via the respiratory route to suppress SARS-CoV-2 infection *in vivo*.

For *in vivo* studies, we used the K18-hACE2 mouse model, in which human ACE2 is expressed under control of the keratin 18 promoter. These mice are exquisitely sensitive to SARS-CoV-2 and succumb to infection with 100% lethality (Zheng et al. 2021; Oladunni et al. 2020; McCray et al. 2007). The experimental approach included a nebulizer delivery apparatus coupled to an animal holder with an integrated nose cone (Fig 4A). This allows for delivery of aerosolized material towards the animal’s breathing space. We refer to this form of ACE2^T371W^ as “aeroACE2^T371W^” to distinguish from the nebACE2^T371W^ generated for cell culture experiments. Mice were treated with buffer alone or aeroACE2^T371W^ 30 minutes or 4 hours prior to infection with 1000 plaque forming units (PFU) SARS-CoV-2 gamma variant. Weight was monitored daily, and mice were euthanized when they reached 80% of starting body weight. In the buffer-treated group, all mice lost weight and were euthanized on day 5 or 6 (Fig 4B-C). Remarkably, in the cohort treated with aeroACE2^T371W^ 30 minutes prior to infection, nearly all mice survived, with only one death on day 7. In a separate cohort, we quantified viral titers in lungs from mice infected for 3 days under the same treatment conditions. Buffer-treated mice had over 7log_10_ PFU per gram tissue, whereas aeroACE2^T371W^-treated mice did not have detectable virus in their lungs at this time point. (Fig S3A). We next pretreated mice with aeroACE2^T371W^ for 4 hours prior to infection to determine if the prophylactic effect would be temporally extended. All mice succumbed to infection, with only a slight delay by 1-2 days in their demise relative to the control group (Fig 4B-C). These data suggest that prophylaxis with aeroACE2^T371W^ relatively soon before SARS-CoV-2 exposure provides strong protection from infection and disease.

**Figure 4.**
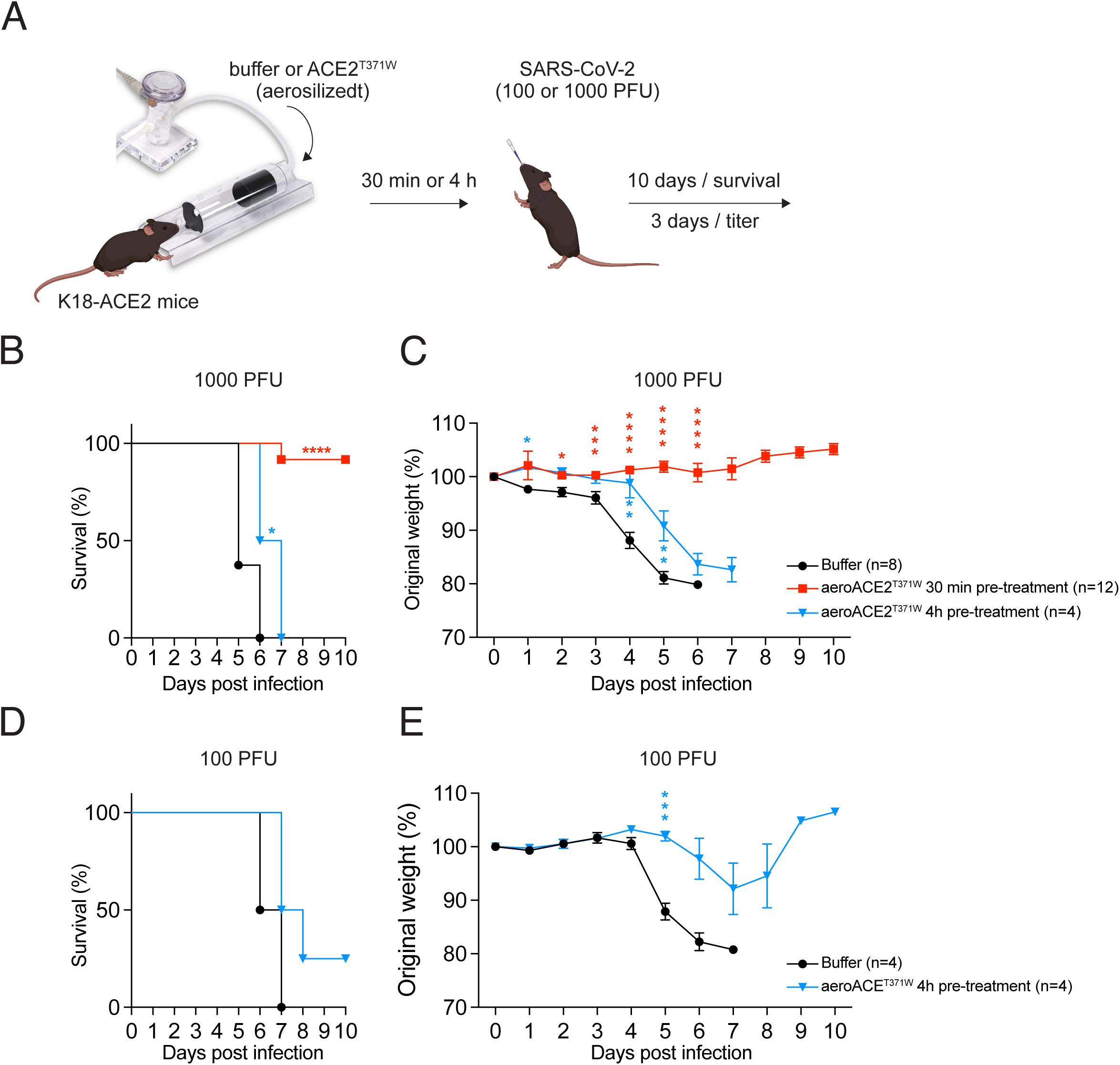
sACE2^T371W^ inhibits SARS-CoV2 replication and increases survival rates in mice. A) Experimental scheme to demonstrate aerosolized treatment and intranasal (IN) infection of K18-hACE2 mice. B) and C) Survival and daily weight of K18-hACE2 mice nebulized with sACE2^T371W^ either 30min prior or 4h prior to Intransal (IN) infection with 1000 CFU P.1 SARS-CoV-2. Data represent means from n= 8 buffer treated mice (2M/6F), n=12 30min prior sACE2^T371W^ treatment (6M/6F), and n=4 4h prior sACE2 treatment (4F) D) and E) Survival and daily weight of K18-hACE2 mice nebulized with sACE2^T371W^ either 30min prior or 4h prior to Intranasal (IN) infection with 100 CFU P.1 SARS-CoV-2. Data represent means of n=4 for each group and all female mice were used. Statistical significance was determined by log-rank (mantel-Cox) tests (B and D) and unpaired t tests (C and E) * p<0.05, ** p<0.01, *** p<0.001, **** p<0.0001.

As the mice treated with aeroACE2^T371W^ 4 hours prior to infection still succumbed, we asked whether a lower viral dose could reveal a protective effect for aeroACE2^T371W^ at this time point. We repeated the experiment at 100 PFU SARS-CoV-2 and found that 100% of control mice and 75% of aeroACE2^T371W^ treated mice succumbed to infection (Fig 4D). While the survival difference was not statistically different given the small cohort, weight loss between the groups was statistically significant (Fig 4E). We extended the prophylactic treatment further by treating mice with aeroACE2^T371W^ 1 day, 2 days, or 1 and 2 days prior to infection. We observed no difference in survival or weight loss relative to untreated mice (Appendix Fig S3B-C). We also treated mice “therapeutically” by administering aeroACE2^T371W^ 30 min after infection. Compared to untreated mice, the treated cohort lost less weight and had a considerable delay in their survival curve (Appendix Fig S3B-C). However, these mice died by day 10, likely due to the extreme sensitivity of the K18-ACE2 model.

## Discussion

In this work, we demonstrate three advances towards using ACE2 as an aerosolized therapeutic for COVID-19. First, we developed an ACE2 variant that enables rapid, efficient purification of the full ACE2 ectodomain without C-terminal proteolysis. Second, we show this protein can be nebulized and recovered intact with no loss of RBD binding and prevents infection by multiple SARS-CoV-2 variants *in vitro*. Finally, we show that aerosolized ACE2^T317W^ prevents disease in an animal model of COVID-19.

The full ACE2 ectodomain has been purified as a secreted protein without an affinity tag using several biochemical separation methods (Vickers et al. 2002). This protein crystallizes via dimeric interactions between symmetry-related monomers that resemble the dimerization seen in the full-length ACE2 dimer (Towler et al. 2004; Yan et al. 2020). In contrast, secreted constructs containing the Enzyme domain but without the Neck domain have been purified for structural studies using standard NiNTA affinity with a C-terminal His tag (Kirchdoerfer et al. 2018; Li 2008; Li et al. 2005; Song et al. 2018; Wu et al. 2012; Lan et al. 2020). We found the full WT ACE2 ectodomain proteolyzes its own C-terminal His_10_ tag, and this His_10_ tag could only be used for purification when the protein was expressed in cells treated with 1 mM EDTA to inhibit metalloprotease activity (Appendix Fig S1A). We have not determined the extent to which WT ACE2 proteolyzes C-terminal portions of the Neck domain in addition to the His tag, but we were eager to preserve this region as it is an important mediator of ACE2 ectodomain dimerization. A soluble dimeric ACE2 construct more faithfully mimics the cell surface receptor, and the avidity effect conferred by dimerization should enhance the ability of soluble ACE2 to compete for virus binding. Proteolytic stability of the C-terminal His_10_ tag ensures purification of soluble ACE2 with the dimerizing Neck domain intact and allows a single-step purification from culture media.

The T371W mutation was designed to project into the substrate binding pocket of ACE2 and prevent substrate binding (Fig 1B). The introduced Trp residue is predicted to occupy the same area as the inhibitor MLN-4760 molecule in the enzyme’s active site (Towler et al. 2004). The T371W mutation does not change the solution behavior of the protein compared to WT ACE2 (Fig 1D), and the mutant ACE2 has the expected dimer MW in solution (Appendix Fig S2B). The T371W mutation largely abolishes the enzymatic activity of ACE2 (Appendix Fig S1), and any residual activity is not inhibited by MLN-4760, confirming disruption of substrate binding by this modification (Appendix Fig S1C). ACE2^T371W^ could be nebulized with >90% recovery and no change in measurable biophysical behavior. The nebulized protein bound the RBDs of SARS-CoV-1 and SARS-CoV-2 equivalently to non-nebulized ACE2^T371W^, with similar affinities to those reported for the WT ACE2 interaction by SPR (Fig 2) (Wrapp et al. 2020). Nebulized or not, our mutant soluble ACE2 inhibited infection by viruses containing an array of spike variants (Fig 3).

The aerosolized ACE2^T371W^ prevented disease when given prophylactically 30 minutes before infection (Fig 4). However, the protein was not as effective when administered at earlier times before viral infection (Appendix Fig S3). We have not measured the rate of clearance of the nebulized ACE2^T371W^ *in vivo*. It is possible that formulations that reduce the rate of clearance of the nebulized ACE2 in the respiratory system would improve its efficacy for longer times prior to exposure. The protein was also not effective when given as a treatment post-infection (Appendix Fig S3). This finding is likely due in part to the nature of the K18-ACE2 mouse model, which over-expresses human ACE2 and thereby highly sensitizes the animals to infection and disease. Further *in vivo* studies will be required to determine whether this protein has higher prophylactic or treatment efficacy in an animal model expressing lower unmodified levels of endogenous ACE2.

Our work demonstrates the utility of an aerosol delivery system to deliver a therapeutic protein and prevent infection in K18-ACE2 mice by SARS-CoV-2. The modified ACE2 described herein is a potential anti-viral agent for any virus that exploits human ACE2 for cellular entry.

## Methods and Protocols

### Cells

HEK-293T cells were maintained in DMEM supplemented with 10% FBS and 1X non-essential amino acids. A549-TMPRSS2-ACE2 cells (a gift from C. Rice, (Hoffmann et al. 2021)) were maintained in DMEM supplemented with 10% FBS, 1X non-essential amino acids, and 0.4 mg/ml Geneticin. Huh7.5 cells were transduced with a lentiviral vector (SCRBBL) expressing a catalytically inactive human ACE2, selected with 10 µg/ml Blasticidin, and maintained in DMEM supplemented with 10% FBS and 1X non-essential amino acids. Spodoptera frugiperda (Sf9) cells were grown in ESF 921 media while shaking at 140 rpm at 27 °C. The cell line was not further authenticated and was not tested for mycoplasma contamination.

### Expression constructs and baculovirus production for protein purification

The pFastBac-ACE2 plasmid was created using Gibson assembly methods to insert a synthetic dsDNA block into pFastBac-1 plasmid that was linearized with BamHI and XhoI. The resulting construct encodes a Melittin signal peptide, the full human ACE2 ectodomain (residues 19-740), a TEV recognition site (ENLYFQG), a TG spacer, and 10 His residues. Mutations were introduced through design in the dsDNA pieces or through standard PCR mutagenesis methods. Mutagenesis primer sequences are available upon request.

Baculovirus were produced in Sf9 cells using the Bac-to-Bac Baculovirus Expression System (ThermoFisher) according to the manufacturer’s instructions.

### Expression and purification of ACE2 ectodomains

Sf9 cells at a density of 2 million cells / mL were infected with 20 mL of P2 baculovirus per Liter of culture and then cultured for 96 hours. Cells were pelleted by centrifugation for 10 min at 6000 x g at 4 °C and the supernatants were either used directly or else supplemented with 20% glycerol and frozen at −80°C until purification. To purify the ACE2 ectodomains, the supernatants were supplemented with protease inhibitors (160 μg/ml benzamidine, 100 μg/ml leupeptin, 1 mM PMSF, and 1 uM E-64), and further clarified by filtration through a 0.2 micron filter. His-tagged ACE2 ectodomains were purified by immobilized metal ion affinity chromatography using Ni-Penta Agarose resin. Supernatants were passed over the resin using gravity. Resin was washed in 50 mM HEPES pH 7.5, 150 mM NaCl, and 20 mM Imidazole pH 7.5. The ACE2 ectodomains were eluted in the same wash buffer supplemented with Imidazole to a final concentration of 300 mM. For animal experiments, the protein was subjected to gel filtration in a superdex 200 column (GE) equilibrated in sterile Phosphate Buffered Saline. Eluted samples were concentrated to ∼10 mg/mL and then supplemented with PEG-8000 to a final concentration of 1% w/v. For experiments other than the peptide cleavage assay, the protein was purified over the same column equilibrated in 20 mM HEPES pH 7.5 and 150 mM NaCl. For the peptide cleavage assay, ACE2 proteins were purified as described above except that 1 mM EDTA was added to the Sf9 cultures at the time of infection and the final gel filtration buffer was 150 mM NaCl, 50 mM HEPES pH 7.5, and 100 µM EDTA.

### SEC-MALS

A Sephadex 200 Increase 10/300 size-exclusion column was equilibrated with gel filtration buffer. 400 µg of the sample at 2 mg / mL was injected and chromatographed at a flow rate of 0.5 mL / min. Data were recorded on a Shimadzu UV detector, a Wyatt TREOS II light-scattering detector, and a Wyatt Optilab t-REX differential refractive-index detector, which were all calibrated with a BSA standard. SEC-MALS data were analyzed with ASTRA version 7.3.0.11.

### Circular Dichroism Spectroscopy

Circular dichroism spectroscopy was performed on MBR’s Jasco J-815 CD Spectrometer. Samples were prepared and placed in a 0.1 cm CD cuvette. As a first step, spectra were collected on the samples from 190 to 250 nm. 208 nm and 222 nm were selected as wavelengths to monitor the CD as a function of temperature, which was varied from 25°C to 95°C at a ramp rate of 2°C/min. Only the data taken at 222 nm were analyzed.

### Isothermal Titration Calorimetry

ITC experiments were carried out in a Malvern iTC200 calorimeter at 20C. a first injection volume of 0.5 μL was employed, followed by 20 injections of 1.9 μL each; all injection rates were 0.5 μL/sec. The stir rate was 750 rpm, and the reference power was 5.0 μcal/s. The time between injections was 120 s, with a data-filtering period of 5 s. ACE2 was included in the cell at a concentration of 11.8 µM and RBD was included in the syringe at the concentration of 120 µM. The data were integrated, analyzed, and illustrated using NITPIC (Keller et al. 2012), SEDPHAT (Houtman et al. 2007), and GUSSI (Brautigam et al. 2016; Brautigam 2015), respectively.

### Thermal stability analysis

Differential Scanning Fluorimetry (DSF) experiments were conducted using a BioRad CFX96 Real-Time System machine. 20 µL reactions were set up with a protein concentration of 5 µM in gel filtration buffer that was supplemented with SYPRO Orange dye at a final concentration of 5x. The temperature was increased from 20°C to 95°C at a rate of ∼1 °C per minute and fluorescence was recorded in all channels. The dye fluoresced most strongly in the HEX channel, and this data was used for all analyses.

### ACE2 Peptide Cleavage Assay

The fluorescent peptide cleavage assay was set up with final concentrations of 0.15 nM protein, 150 mM NaCl, 50 mM HEPES, 0.01% Nonidet P-40 and 1 µM EDTA. Where indicated, ZnCl_2_ and MLN-4760 were added to a final concentration of 10 µM. The protein was incubated in these conditions at RT for 30 min and then the reaction was initiated by the addition of Mca-APK(Dnp)-OH trifluoroacetate to a final concentration of 50 µM. The reaction was analyzed at RT at the indicated time points using a CLARIOstar plate reader (BMG LABTECH) with excitation at 320 nm and emission at 405 nm.

### Immunoblot analysis

Samples were mixed with 5x SDS-PAGE loading buffer, heated at 37 °C for 5 min, and subjected to 10% SDS-PAGE. Proteins were transferred to nitrocellulose filters using Bio-Rad Trans Blot Turbo system and immunoblotted with a 1:1000 dilution of either monoclonal anti-His clone His.H8 or anti-ACE2 (Abcam ab108252). Bound antibodies were visualized by chemiluminescence (SuperSignal West Pico Chemiluminescent Substrate, Thermo Scientific, Waltham, MA) using a 1:5000 dilution of anti-mouse IgG or anti-rabbit IgG conjugated to horseradish peroxidase (Jackson ImmunoResearch Laboratories, Inc., West Grove, PA). The filters were exposed to Blue X-ray Film (Phoenix Research Products, Pleasanton, CA).

#### Viruses

Virus from infectious clone pCC1-4K-SARS-CoV-2-Wuhan-Hu-1-ZsGreen (a gift from S. Wilson) was generated and propagated as previously described (Mar et al. 2023). SARS-CoV-2, isolate hCoV-19/USA/MD-HP05647/2021 (Lineage B.1.617.2; Delta Variant) (BEI) was propagated as previously described for other SARS-CoV-2 variants (Mar et al. 2023). SARS-CoV-2 (hCoV-19/Japan/TY7-503/2021, P.1) was obtained from BEI Resources and inoculated onto Vero-E6-C1008 (ATCC, VA) plated at a density of 7 x 10^6^ cells in a 175 cm^2^ tissue culture flask in 2% FBS (Gibco, MA)/1X non-essential amino acids (Gibco, MA)/MEM (Gibco, MA) for 45 minutes at 37°C. Viral inoculum was removed and replaced with 2% FBS (Gibco, MA)/1X non-essential amino acids (Gibco, MA)/MEM (Gibco, MA) and cells were incubated for 3 days at 37°C before supernatants were clarified by centrifugation at 1000 x g for 5 minutes before aliquoting and storage at −80°C. Viral titer was determined by plaque assay on Vero-E6-C1008 cells (ATCC, VA). In brief, cells were plated at 6.5 x 10^5^ cells per well of a 6 well plate and infected with 6 dilutions from a 10-fold dilution series in 1% FBS (Gibco, MA)/1X non-essential amino acids (Gibco, MA)/MEM (Gibco, MA) for 30 minutes at 37°C. Inoculum was then removed and replaced with Avicel overlay medium (5% FBS [Gibco, MA]/1X penicillin-streptomycin [Gibco, MA]/1X GlutaMAX [Gibco, MA]/1X Modified Eagle Medium, Temin’s Modification #11935-046 [Gibco, MA]/1.2% Avicel RC-591 [DuPont, DE]). Cells were incubated for 3 days at 37°C before overlay medium was removed and cells were fixed with 4% PFA (Sigma-Aldrich, MA) for 30 minutes at room temperature. Fixative was then removed, and cells were stained for at least 1 hr with crystal violet (0.2% crystal violet [Sigma-Aldrich, MA]/20% ethanol [Sigma-Aldrich, MA]) before removal and plaque enumeration. Human coronavirus OC43 (ATCC strain VR-1558) was propagated as previously described (Boys et al. 2020). Lentivirus and OC43 infections were performed in a Biosafety Level 2 (BSL2) facility and SARS-CoV-2 infections were performed in a Biosafety Level 3 (BSL3) facility, each equipped with the appropriate safety features to prevent bio-hazardous exposure or unintentional pathogen release. Experiments and biosafety protocols were reviewed and approved by the Institutional Biosafety Committee, according to guidelines provided by the UT Southwestern Office of Business and Safety Continuity.

#### Lentiviral Pseudoparticle Production

Spike-pseudotyped HIV-1-based lentiviral vectors were generated by co-transfecting 293T cells with expression plasmids pSCRBBL-Fluc (Richardson et al. 2018), pLV-Spike (G614 variant, Alpha B.1.1.7 variant, Beta B.1.351 variant, Gamma P.1 variant, Delta B.1.617.2 variant, Omicron BA.4 & BA.5 variant) (Invivogen), and pCMV.gag-pol using XtremeGene9. Six hours post-transfection, media was replaced with DMEM containing 3% FBS and 1X non-essential amino acids. Supernatants were collected at 48 and 72 hrs, pooled, supplemented with 20mM HEPES and 4 ug/ml Polybrene, and aliquoted.

Control lentiviruses were generated the same way but pseudotyped with VSV glycoprotein using plasmid pCMV.VSVg.

#### 10 dilution drug curves and transduction with SARS-CoV-2 pseudotyped lentivirus

A 3-fold dilution series of sACE2 starting at 333µM, was diluted in media containing DMEM, 3% FBS, 1X NEAA, 20 mM HEPES, and 4 µg/ml Polybrene. An equal volume of each sACE2 dilution was incubated with an equal volume of each spike-pseudotyped lentivirus or VSVg-pseudoyped control for 30 minutes at 37°C. 10,000 Huh7.5mutACE2 cells were added to each well and incubated for 48 hrs. Media was aspirated and 20 ul of 1X passive lysis buffer was added to each well, mixed on orbital shaker at 900 rpm for 1 min, and stored at −80°C. Luciferase activity was detected by Luciferase Assay System and quantified on a Berthold Centro XS^3^ LB 960 luminometer.

#### ACE2 inhibitory studies in cell culture

A549-TMPRSS2-ACE2 cells were plated at a density of 80,000-100,000 c/well. A 3-fold dilution series of sACE2^T371W^ or nebACE2^T371W^, starting at 1µM, was diluted in infection media containing 1% FBS and 1X NEAA. An equal volume of each dilution was incubated with an equal volume of hOC43, SARS-CoV-2-zsGreen, or SARS-CoV-2-delta for 30 minutes at 33°C (OC43) or 37°C (SARS-CoV-2). Media was aspirated from cells and ACE2/virus complex was added to cells and incubated for 30-60 minutes, followed by addition of complete media containing 10%FBS and 1X NEAA. Cells were incubated for 7-24 hrs. Cells were fixed with a final concentration of 1 or 4% PFA and run on flow cytometer or antibody stained followed by flow cytometry.

#### Antibody staining

After initial fixation, cells were incubated with BD Cytofix solution and permeabilized with BD Cytoperm solution (BD Cytofix/Cytoperm Kit). Cells infected with OC43 were stained with anti-Coronavirus Group Antigen Antibody, nucleoprotein of OC-43, clone 542-7D (Millipore) at a dilution of 1:500 and goat anti-mouse Alexa Fluor 488 secondary at a dilution of 1:1000. Cells infected with SARS-CoV-2 delta variant were stained with SARS-CoV/SARS-CoV-2 Nucleocapsid Antibody at a dilution of 1:1000 (Sino Biological) and goat anti-mouse Alexa Fluor 488 secondary at a dilution of 1:1000 (Life Technologies). Stained cells were analyzed on a Stratedigm S1000EON bench flow cytometer.

#### Mice

Male and female K18-hACE2 mice (Jackson Laboratory) 6-8 weeks were used for these studies. Mice were housed according to NIH guidelines for housing and care in a Biosafety Level 3 animal laboratory. All procedures were approved by the Institutional Animal Care and Use Committee (protocol number 2020-102987) of University of Texas Southwestern.

#### In Vivo Infections and Intranasal and Nebulizer Treatments

K18-hACE2 mice were anesthetized with ketamine/xylazine (80/6 mg/kg) and treated with soluble ACE2^T371W^ (sACE2 ^T371W^; 8.5-12 mg/ml) or buffer control using a nebulizer (Animal holder nebulizer delivery system, Kent Scientific). Each mouse was placed in the holder and 125ul of sACE2^T371W^ or buffer was added to the nebulizer and nebulized at 0.2L/min. After the vaper was cleared, each mouse was removed from the restraint and the process was repeated for each additional mouse. At 30 min post-treatment, mice were intranasally (IN) infected with 1 x 10^3^ PFU of SARS-CoV-2 P.1 variant suspended in 30 µl of phosphate-buffered saline (PBS). For low dose infections, K18-hACE2 mice were anesthetized with ketamine/xylazine (80/6 mg/kg) and intranasally (IN) infected with 1 x 10^2^ PFU of SARS-CoV-2 P.1 variant suspended in 30 µl of phosphate-buffered saline (PBS) and treated 20 µl IN with soluble ACE2^T371W^ (sACE2^T371W^; 11.6-12 mg/ml) or control 30 min either before or after infection or given 1dpi, 2dpi, or combination of 1dpi and 2dpi. All mice were monitored daily for weight and mortality. Animals that lost more than 20% of their original body weight or appear lethargic, hunched, or unable to obtain food were euthanized per Institutional Animal Care and Use Committee guidelines.

For quantifying SARS-CoV-2 viral load, whole lungs were homogenized in 600ul 2% FBS (Gibco)/MEM (Gibco) with stainless steel beads (NextAdvance) in a Bullet Blender Pro (NextAdvance). Lung tissue was weighed before homogenization. Tissue homogenates were clarified of debris by centrifugation at 4°C at 800g for 5min, and aliquots for plaque assay were frozen at −80°C. Plaque assays were performed as described above.

## Structured Methods

### Reagents and Tools Table

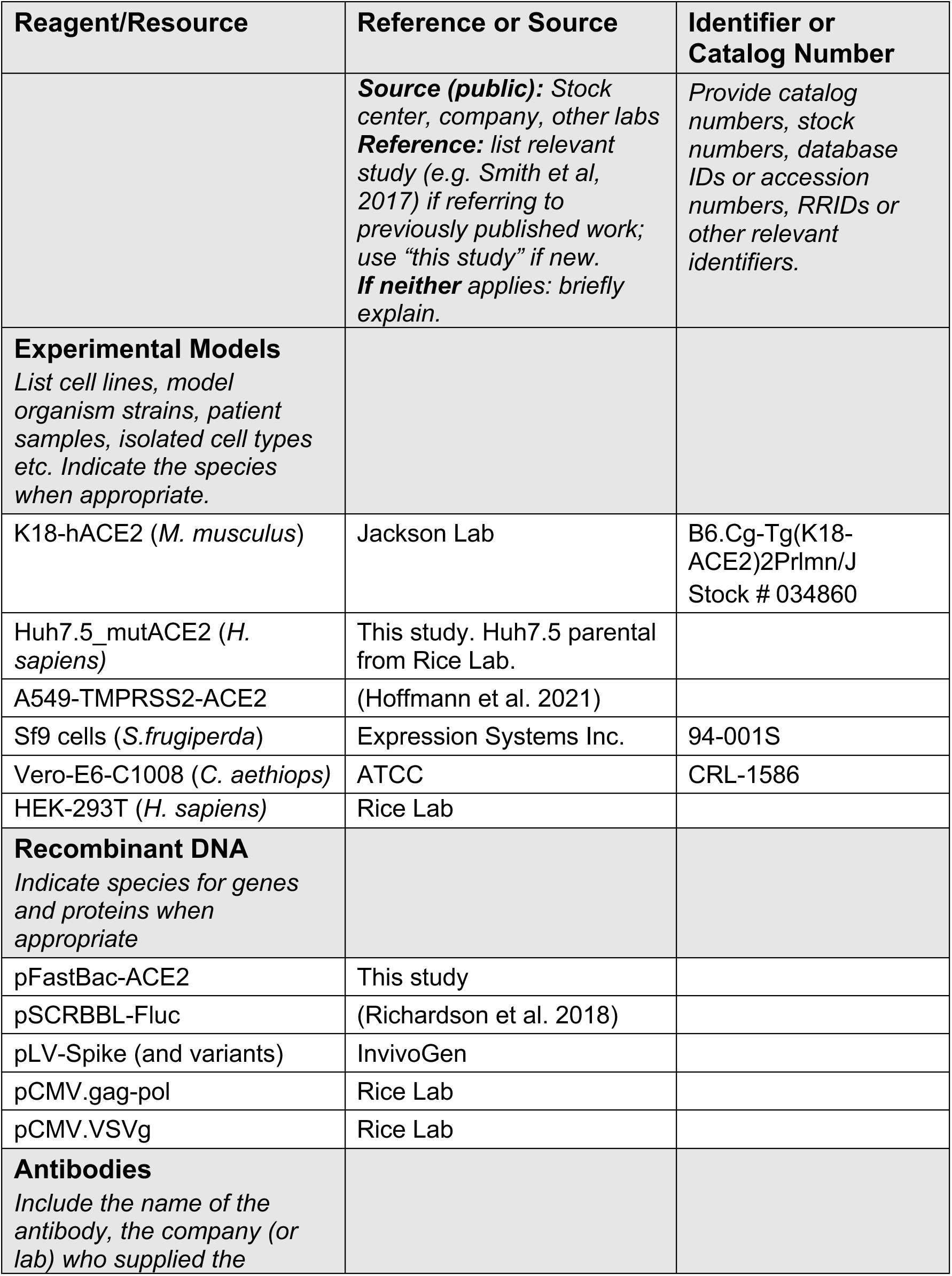
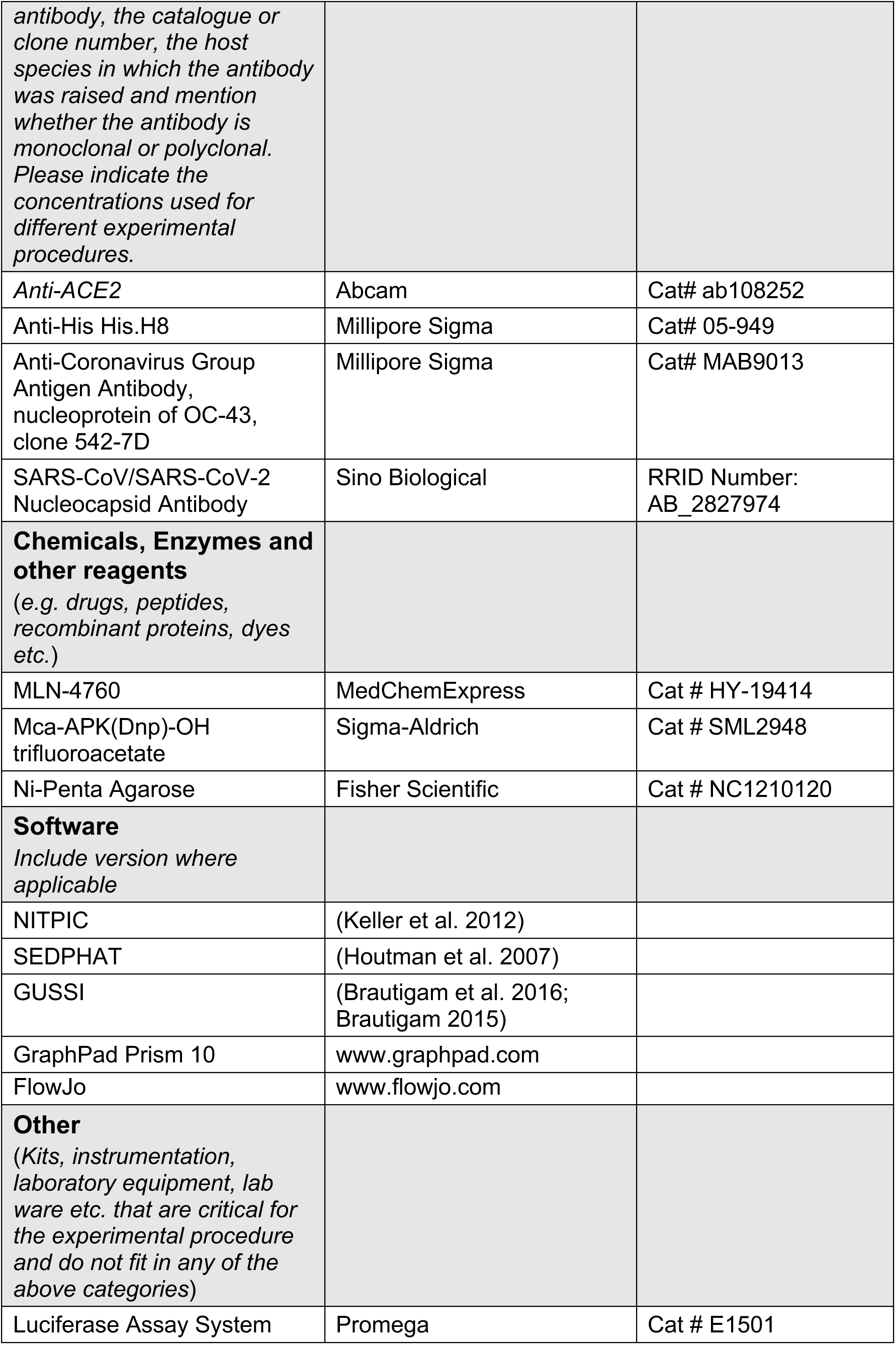
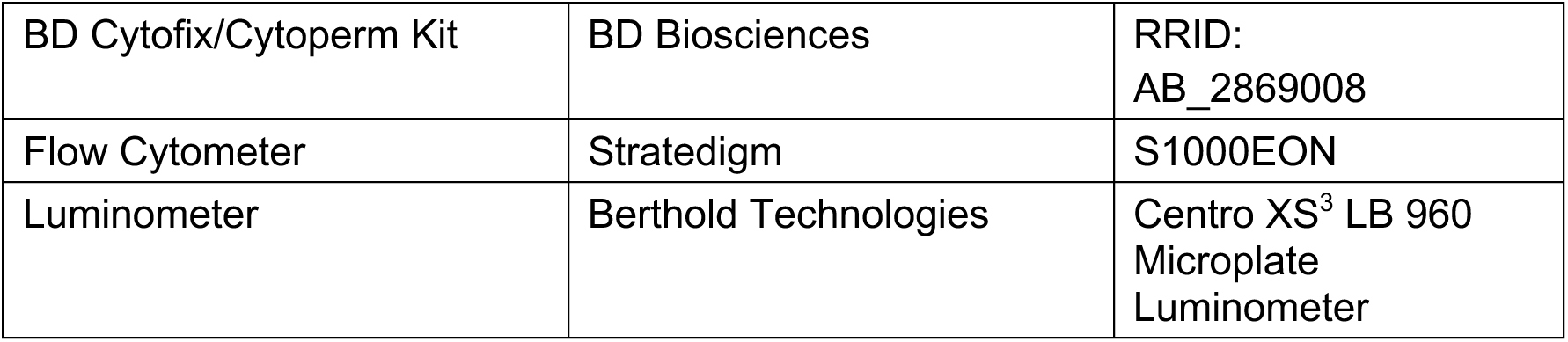

## Acknowledgements

We thank Gabriel Glenn Gregorio for assistance with purifying ACE2^T371W^. We thank Chad Brautigam and Shih-Chia Tso for carrying out CD, SEC-MALS, and ITC experiments.

This project was supported in part by the NIH 4R00GM141261 (to DLK) and R35GM116387 (to DMR), a Welch Foundation grant (I-1770) to DMR and a National Science Foundation’s Small Business Innovation Research (NSF SBIR) grant, awarded to White Rock Therapeutics, with a subcontract to J.W.S. at UT Southwestern (Grant No. 2136508). J.W.S. is a Burroughs Wellcome Fund Investigator in the Pathogenesis of Infectious Disease.

## Abstract Importance

The rapid evolution of SARS-CoV-2 variants poses a challenge for immune recognition and antibody therapies. However, the virus is constrained by the requirement that it recognizes a human host receptor protein. A recombinant ACE2 could protect against SARS-CoV-2 infection by functioning as a soluble decoy receptor. We designed a mutant version of ACE2 with impaired catalytic activity to enable purification of the protein using a single affinity purification step. This protein can be nebulized and retains the ability to bind the relevant domains from SARS-CoV-1 and SARS-CoV-2. Moreover, this protein inhibits viral infection against a panel of coronaviruses in cells. Finally, we developed an aerosolized delivery system for animal studies and show the modified ACE2 offers protection in an animal model of COVID-19. These results show proof-of-concept for an aerosolized delivery method to evaluate anti-SARS-CoV-2 agents *in vivo* and suggest a new tool in the ongoing fight against SARS-CoV-2.

---

**Figure S1.**
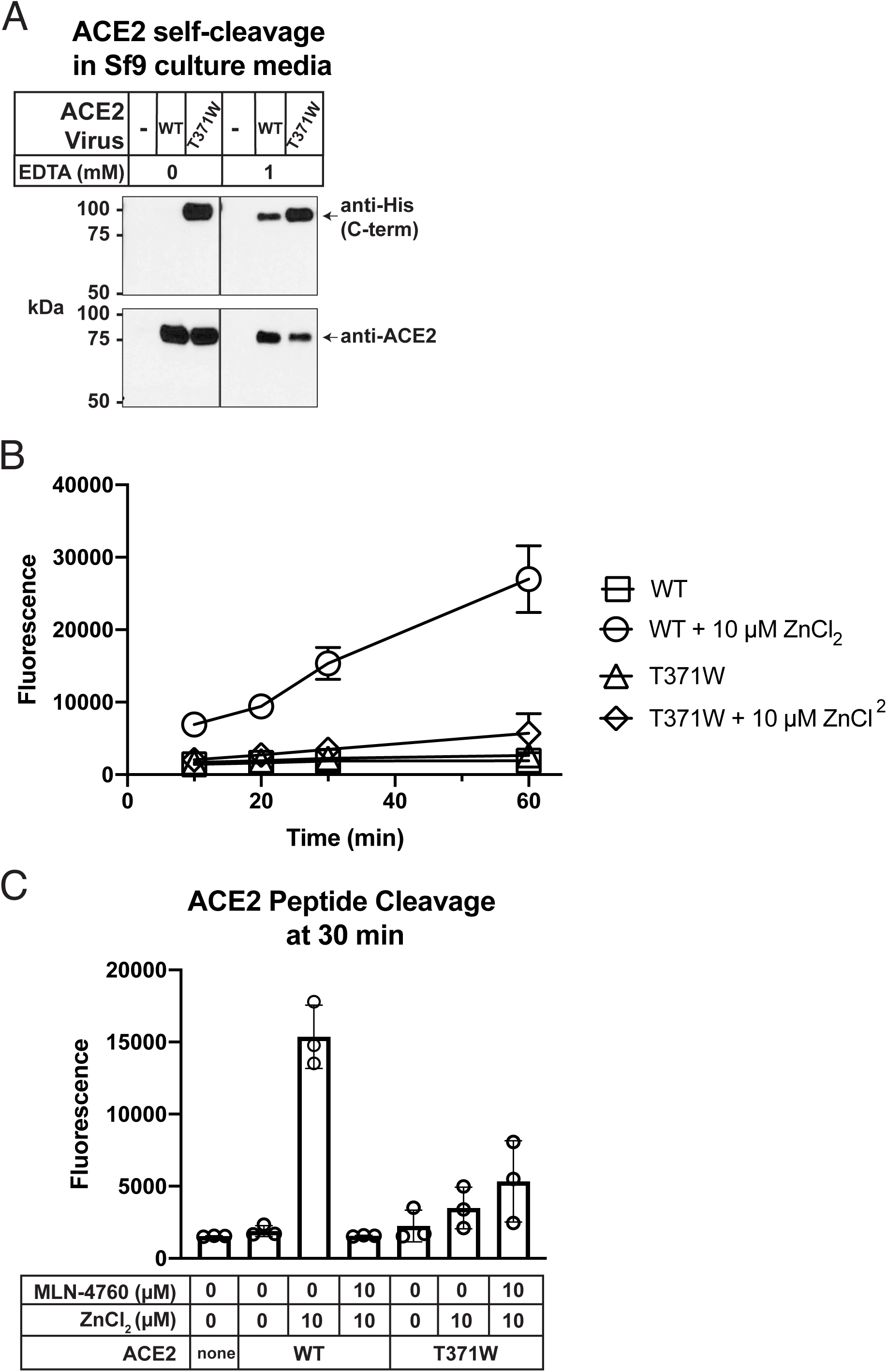
Characterization of ACE2^T371W^ enzymatic activity. A) ACE2 self-cleavage in insect cell culture media. Sf9 cells were infected with the indicated virus for 48 hrs. Conditioned supernatants were collected and subjected to immunoblot analysis with either anti-ACE2 or anti-His antibodies. B) Peptide cleavage by ACE2 ectodomains monitored through Mca fluorescence. Proteins were treated with ZnCl_2_ as indicated and fluorescence was recorded at the indicated time points. C) ACE2 peptide cleavage at 30 minutes monitored through Mca fluorescence in the presence or absence of ZnCl2 and MLN-4760 inhibitor, as indicated.

**Figure S2.**
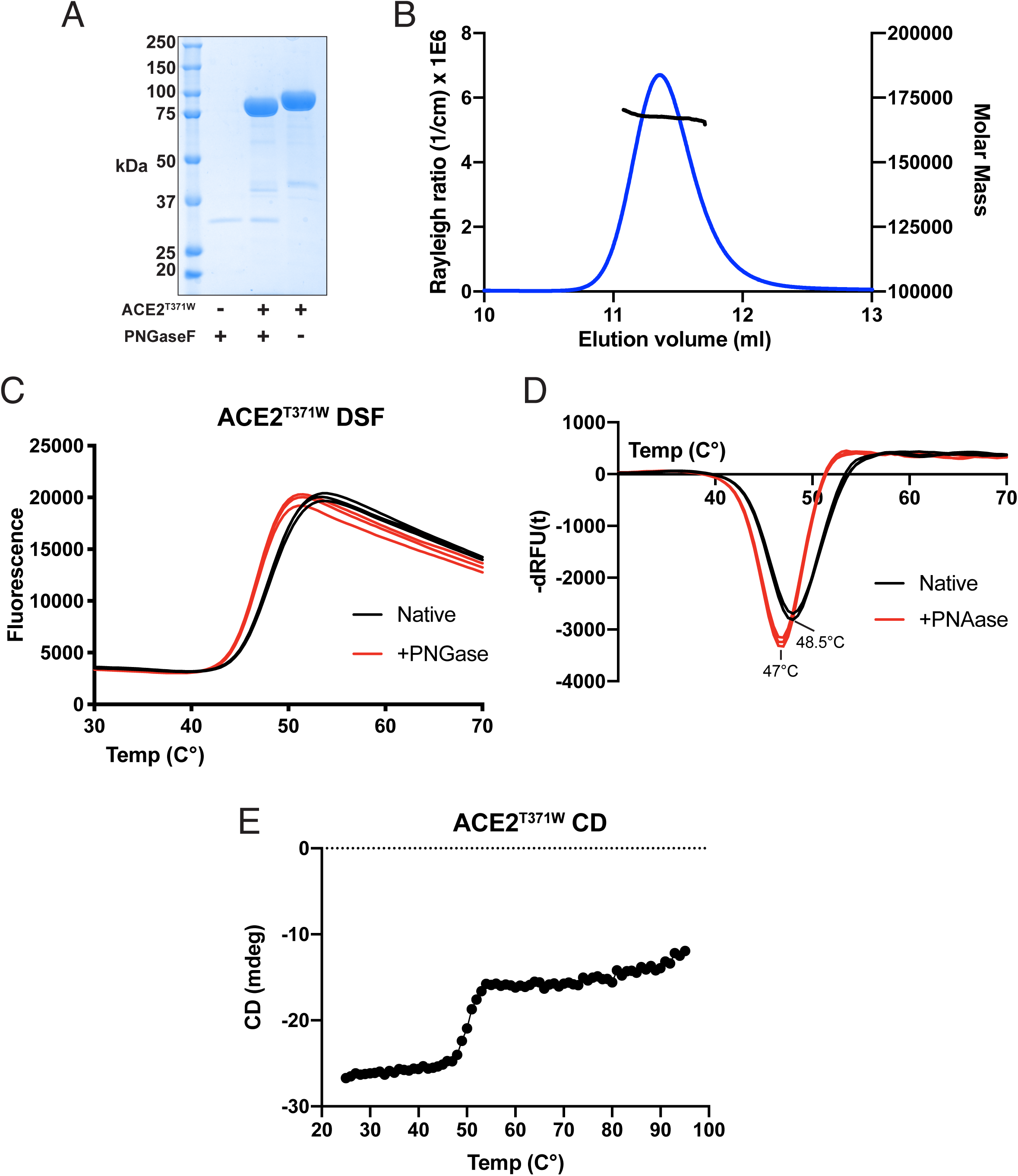
Solution characterization of ACE2^T371W^ ectodomains. A) Coomassie-stained SDS-PAGE analysis of ACE2^T371W^ ectodomains subjected to PNGaseF treatment. B) SEC-MALS analysis of purified ACE2^T371W^ ectodomains. Rayleigh scattering is shown in blue trace and the calculated molar mass is show in the black trace. C) Fluorescent traces from DSF experiment with native and PNGaseF-digested _ACE2T371W._ D) Derivative curves from the fluorescent traces shown in (C). E) CD melt trace for signal at 222 nm for ACE2^T371W^.

**Figure S3.**
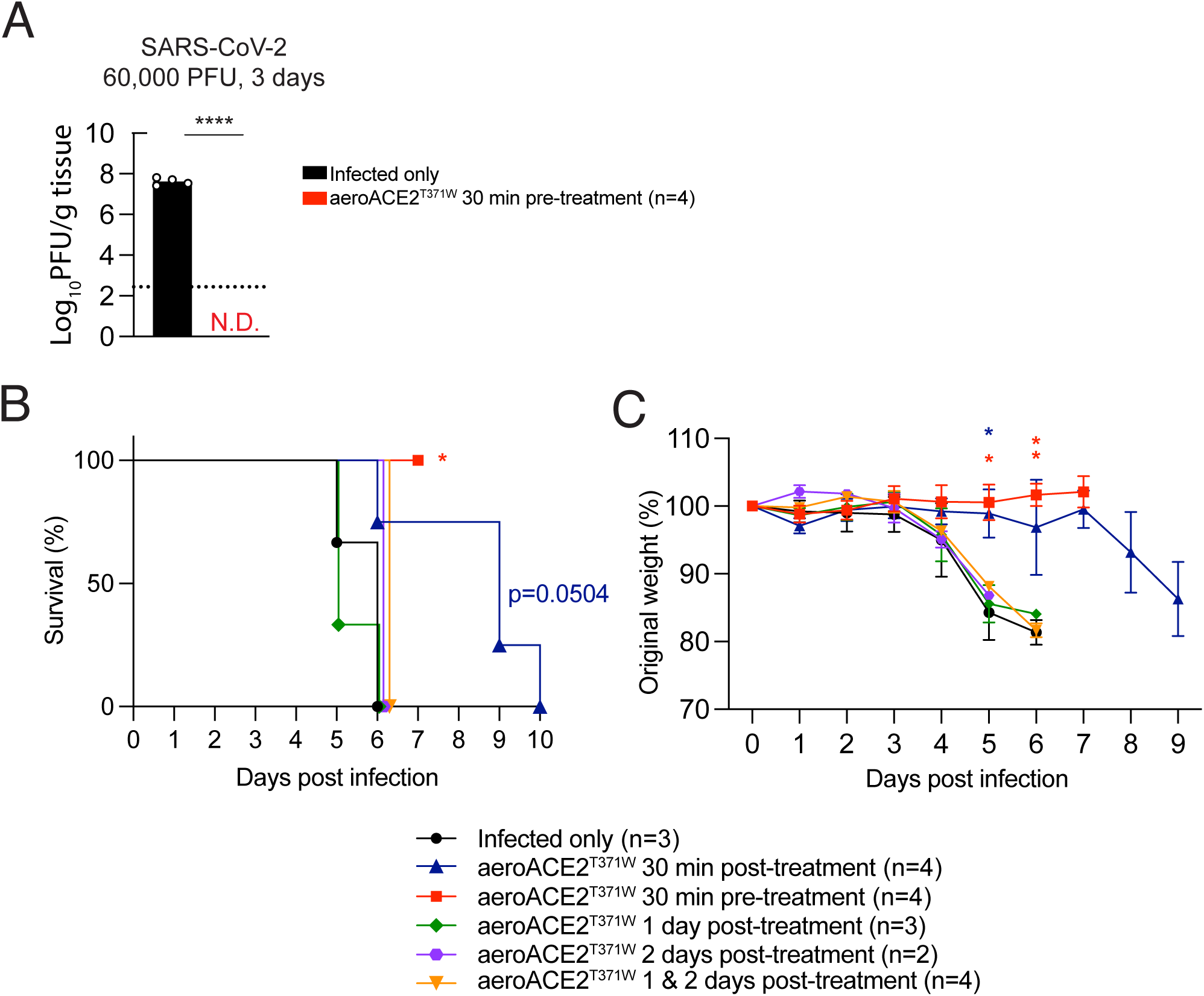
Intranasal pretreatment of sACE2 inhibits SARS-CoV-2 replication. A) K18-hACE2 mice were treated intranasally with sACE2^T371W^ 30min prior to infection or infected alone with 60,000 PFU SARS-CoV-2 P.1 and assessed for viral burden in the lungs by plaque assay at 3 days post infection. Data represent means of n= 4 K18-hACE2. All male mice were used for this experiment. B) and C) Survival and daily weight of K18-hACE2 mice treated intranasally either 30min prior to infection with SARS-CoV-2 P.1 or 30min, 1 day, 2 day, or 1+2 day post infection or infected alone. Infected only and 30min prior to infection treatment were given 60,000 PFU of SARS-CoV-2 P.1. All other groups were infected with 100 PFU of SARS-CoV-2 P.1. All female mice were used for these experiments. Statistical significance was determined by log-rank (mantel-Cox) tests (B) and multiple unpaired t tests (C) * p<0.05 and ** p<0.01.

## References Cited

Boys, Ian N., Elaine Xu, Katrina B. Mar, Pamela C. De La Cruz-Rivera, Jennifer L. Eitson, Benjamin Moon, and John W. Schoggins. 2020. ‘RTP4 Is a Potent IFN-Inducible Anti-flavivirus Effector Engaged in a Host-Virus Arms Race in Bats and Other Mammals’, Cell host & microbe, 28: 712–23.e9.

Brautigam, Chad A. 2015. ’Chapter Five - Calculations and Publication-Quality Illustrations for Analytical Ultracentrifugation Data.’ in James L. Cole (ed.), Methods in Enzymology (Academic Press).

Brautigam, Chad A., Huaying Zhao, Carolyn Vargas, Sandro Keller, and Peter Schuck. 2016. ‘Integration and global analysis of isothermal titration calorimetry data for studying macromolecular interactions’, Nature protocols, 11: 882–94.

Cele, Sandile, Laurelle Jackson, David S. Khoury, Khadija Khan, Thandeka Moyo-Gwete, Houriiyah Tegally, James Emmanuel San, Deborah Cromer, Cathrine Scheepers, Daniel G. Amoako, Farina Karim, Mallory Bernstein, Gila Lustig, Derseree Archary, Muneerah Smith, Yashica Ganga, Zesuliwe Jule, Kajal Reedoy, Shi-Hsia Hwa, Jennifer Giandhari, Jonathan M. Blackburn, Bernadett I. Gosnell, Salim S. Abdool Karim, Willem Hanekom, Mary-Ann Davies, Marvin Hsiao, Darren Martin, Koleka Mlisana, Constantinos Kurt Wibmer, Carolyn Williamson, Denis York, Rohen Harrichandparsad, Kobus Herbst, Prakash Jeena, Thandeka Khoza, Henrik Kløverpris, Alasdair Leslie, Rajhmun Madansein, Nombulelo Magula, Nithendra Manickchund, Mohlopheni Marakalala, Matilda Mazibuko, Mosa Moshabela, Ntombifuthi Mthabela, Kogie Naidoo, Zaza Ndhlovu, Thumbi Ndung’u, Nokuthula Ngcobo, Kennedy Nyamande, Vinod Patel, Theresa Smit, Adrie Steyn, Emily Wong, Anne von Gottberg, Jinal N. Bhiman, Richard J. Lessells, Mahomed-Yunus S. Moosa, Miles P. Davenport, Tulio de Oliveira, Penny L. Moore, Alex Sigal, S. A. Ngs, and Commit-Kzn Team. 2022. ‘Omicron extensively but incompletely escapes Pfizer BNT162b2 neutralization’, Nature, 602: 654–56.

Donoghue, Mary, Frank Hsieh, Elizabeth Baronas, Kevin Godbout, Michael Gosselin, Nancy Stagliano, Michael Donovan, Betty Woolf, Keith Robison, Raju Jeyaseelan, Roger E. Breitbart, and Susan Acton. 2000. ‘A Novel Angiotensin-Converting Enzyme–Related Carboxypeptidase (ACE2) Converts Angiotensin I to Angiotensin 1-9’, Circulation Research, 87: e1–e9.

Haschke, Manuel, Manfred Schuster, Marko Poglitsch, Hans Loibner, Marc Salzberg, Marcel Bruggisser, Joseph Penninger, and Stephan Krähenbühl. 2013. ‘Pharmacokinetics and Pharmacodynamics of Recombinant Human Angiotensin-Converting Enzyme 2 in Healthy Human Subjects’, Clinical Pharmacokinetics, 52: 783–92.

Hoffmann, H. Heinrich, Francisco J. Sánchez-Rivera, William M. Schneider, Joseph M. Luna, Yadira M. Soto-Feliciano, Alison W. Ashbrook, Jérémie Le Pen, Andrew A. Leal, Inna Ricardo-Lax, Eleftherios Michailidis, Yuan Hao, Ansgar F. Stenzel, Avery Peace, Johannes Zuber, C. David Allis, Scott W. Lowe, Margaret R. MacDonald, John T. Poirier, and Charles M. Rice. 2021. ‘Functional interrogation of a SARS-CoV-2 host protein interactome identifies unique and shared coronavirus host factors’, Cell host & microbe, 29: 267–80.e5.

Hoffmann, Markus, Hannah Kleine-Weber, Simon Schroeder, Nadine Krüger, Tanja Herrler, Sandra Erichsen, Tobias S. Schiergens, Georg Herrler, Nai-Huei Wu, Andreas Nitsche, Marcel A. Müller, Christian Drosten, and Stefan Pöhlmann. 2020. ‘SARS-CoV-2 Cell Entry Depends on ACE2 and TMPRSS2 and Is Blocked by a Clinically Proven Protease Inhibitor’, Cell, 181: 271–80.e8.

Hoffmann, Markus, Nadine Krüger, Sebastian Schulz, Anne Cossmann, Cheila Rocha, Amy Kempf, Inga Nehlmeier, Luise Graichen, Anna-Sophie Moldenhauer, Martin S. Winkler, Martin Lier, Alexandra Dopfer-Jablonka, Hans-Martin Jäck, Georg M. N. Behrens, and Stefan Pöhlmann. 2022. ‘The Omicron variant is highly resistant against antibody-mediated neutralization: Implications for control of the COVID-19 pandemic’, Cell, 185: 447–56.e11.

Houtman, Jon C. D., Patrick H. Brown, Brent Bowden, Hiroshi Yamaguchi, Ettore Appella, Lawrence E. Samelson, and Peter Schuck. 2007. ‘Studying multisite binary and ternary protein interactions by global analysis of isothermal titration calorimetry data in SEDPHAT: Application to adaptor protein complexes in cell signaling’, Protein Science, 16: 30–42.

Keller, Sandro, Carolyn Vargas, Huaying Zhao, Grzegorz Piszczek, Chad A. Brautigam, and Peter Schuck. 2012. ‘High-Precision Isothermal Titration Calorimetry with Automated Peak-Shape Analysis’, Analytical Chemistry, 84: 5066–73.

Kirchdoerfer, Robert N., Nianshuang Wang, Jesper Pallesen, Daniel Wrapp, Hannah L. Turner, Christopher A. Cottrell, Kizzmekia S. Corbett, Barney S. Graham, Jason S. McLellan, and Andrew B. Ward. 2018. ‘Stabilized coronavirus spikes are resistant to conformational changes induced by receptor recognition or proteolysis’, Scientific Reports, 8: 15701.

Lan, Jun, Jiwan Ge, Jinfang Yu, Sisi Shan, Huan Zhou, Shilong Fan, Qi Zhang, Xuanling Shi, Qisheng Wang, Linqi Zhang, and Xinquan Wang. 2020. ‘Structure of the SARS-CoV-2 spike receptor-binding domain bound to the ACE2 receptor’, Nature.

Li, Fang. 2008. ‘Structural Analysis of Major Species Barriers between Humans and Palm Civets for Severe Acute Respiratory Syndrome Coronavirus Infections’, Journal of virology, 82: 6984–91.

Li, Fang, Wenhui Li, Michael Farzan, and Stephen C. Harrison. 2005. ‘Structure of SARS Coronavirus Spike Receptor-Binding Domain Complexed with Receptor’, Science, 309: 1864–68.

Li, Wenhui, Michael J. Moore, Natalya Vasilieva, Jianhua Sui, Swee Kee Wong, Michael A. Berne, Mohan Somasundaran, John L. Sullivan, Katherine Luzuriaga, Thomas C. Greenough, Hyeryun Choe, and Michael Farzan. 2003. ‘Angiotensin-converting enzyme 2 is a functional receptor for the SARS coronavirus’, Nature, 426: 450–54.

Liu, Lihong, Sho Iketani, Yicheng Guo, Jasper F. W. Chan, Maple Wang, Liyuan Liu, Yang Luo, Hin Chu, Yiming Huang, Manoj S. Nair, Jian Yu, Kenn K. H. Chik, Terrence T. T. Yuen, Chaemin Yoon, Kelvin K. W. To, Honglin Chen, Michael T. Yin, Magdalena E. Sobieszczyk, Yaoxing Huang, Harris H. Wang, Zizhang Sheng, Kwok-Yung Yuen, and David D. Ho. 2022. ‘Striking antibody evasion manifested by the Omicron variant of SARS-CoV-2’, Nature, 602: 676–81.

Mar, Katrina B., Alexandra I. Wells, Marley C. Caballero Van Dyke, Alexandra H. Lopez, Jennifer L. Eitson, Wenchun Fan, Natasha W. Hanners, Bret M. Evers, John M. Shelton, and John W. Schoggins. 2023. ‘LY6E is a pan-coronavirus restriction factor in the respiratory tract’, Nature Microbiology, 8: 1587–99.

McCray, Paul B., Lecia Pewe, Christine Wohlford-Lenane, Melissa Hickey, Lori Manzel, Lei Shi, Jason Netland, Hong Peng Jia, Carmen Halabi, Curt D. Sigmund, David K. Meyerholz, Patricia Kirby, Dwight C. Look, and Stanley Perlman. 2007. ’Lethal Infection of K18-*hACE2* Mice Infected with Severe Acute Respiratory Syndrome Coronavirus’, Journal of virology, 81: 813–21.

Monteil, Vanessa, Matheus Dyczynski, Volker M Lauschke, Hyesoo Kwon, Gerald Wirnsberger, Sonia Youhanna, Haibo Zhang, Arthur S Slutsky, Carmen Hurtado del Pozo, Moritz Horn, Nuria Montserrat, Josef M Penninger, and Ali Mirazimi. 2021. ’Human soluble ACE2 improves the effect of remdesivir in SARS-CoV-2 infection’, EMBO Molecular Medicine, 13: e13426.

Oladunni, Fatai S., Jun-Gyu Park, Paula A. Pino, Olga Gonzalez, Anwari Akhter, Anna Allué-Guardia, Angélica Olmo-Fontánez, Shalini Gautam, Andreu Garcia-Vilanova, Chengjin Ye, Kevin Chiem, Colwyn Headley, Varun Dwivedi, Laura M. Parodi, Kendra J. Alfson, Hilary M. Staples, Alyssa Schami, Juan I. Garcia, Alison Whigham, Roy Neal Platt, Michal Gazi, Jesse Martinez, Colin Chuba, Stephanie Earley, Oscar H. Rodriguez, Stephanie Davis Mdaki, Katrina N. Kavelish, Renee Escalona, Cory R. A. Hallam, Corbett Christie, Jean L. Patterson, Tim J. C. Anderson, Ricardo Carrion, Edward J. Dick, Shannan Hall-Ursone, Larry S. Schlesinger, Xavier Alvarez, Deepak Kaushal, Luis D. Giavedoni, Joanne Turner, Luis Martinez-Sobrido, and Jordi B. Torrelles. 2020. ’Lethality of SARS-CoV-2 infection in K18 human angiotensin-converting enzyme 2 transgenic mice’, Nature Communications, 11: 6122.

Richardson, R. Blake, Maikke B. Ohlson, Jennifer L. Eitson, Ashwani Kumar, Matthew B. McDougal, Ian N. Boys, Katrina B. Mar, Pamela C. De La Cruz-Rivera, Connor Douglas, Genevieve Konopka, Chao Xing, and John W. Schoggins. 2018. ‘A CRISPR screen identifies IFI6 as an ER-resident interferon effector that blocks flavivirus replication’, Nature Microbiology, 3: 1214–23.

Shoemaker, Robert H., Reynold A. Panettieri, Jr., Steven K. Libutti, Howard S. Hochster, Norman R. Watts, Paul T. Wingfield, Philipp Starkl, Lisabeth Pimenov, Riem Gawish, Anastasiya Hladik, Sylvia Knapp, Daniel Boring, Jonathan M. White, Quentin Lawrence, Jeremy Boone, Jason D. Marshall, Rebecca L. Matthews, Brian D. Cholewa, Jeffrey W. Richig, Ben T. Chen, David L. McCormick, Romana Gugensberger, Sonja Höller, Josef M. Penninger, and Gerald Wirnsberger. 2022. ‘Development of an aerosol intervention for COVID-19 disease: Tolerability of soluble ACE2 (APN01) administered via nebulizer’, PloS one, 17: e0271066.

Song, Wenfei, Miao Gui, Xinquan Wang, and Ye Xiang. 2018. ‘Cryo-EM structure of the SARS coronavirus spike glycoprotein in complex with its host cell receptor ACE2’, PLoS pathogens, 14: e1007236.

Tipnis, Sarah R., Nigel M. Hooper, Ralph Hyde, Eric Karran, Gary Christie, and Anthony J. Turner. 2000. ‘A Human Homolog of Angiotensin-converting Enzyme: CLONING AND FUNCTIONAL EXPRESSION AS A CAPTOPRIL-INSENSITIVE CARBOXYPEPTIDASE*’, Journal of Biological Chemistry, 275: 33238–43.

Towler, Paul, Bart Staker, Sridhar G. Prasad, Saurabh Menon, Jin Tang, Thomas Parsons, Dominic Ryan, Martin Fisher, David Williams, Natalie A. Dales, Michael A. Patane, and Michael W. Pantoliano. 2004. ‘ACE2 X-Ray Structures Reveal a Large Hinge-bending Motion Important for Inhibitor Binding and Catalysis’, Journal of Biological Chemistry, 279: 17996–8007.

Turner, Anthony. 2015. ‘ACE2 Cell Biology, Regulation, and Physiological Functions’, The Protective Arm of the Renin Angiotensin System (RAS): Functional Aspects and Therapeutic Implications: 185–89.

Vickers, Chad, Paul Hales, Virendar Kaushik, Larry Dick, James Gavin, Jin Tang, Kevin Godbout, Thomas Parsons, Elizabeth Baronas, Frank Hsieh, Susan Acton, Michael Patane, Andrew Nichols, and Peter Tummino. 2002. ‘Hydrolysis of Biological Peptides by Human Angiotensin-converting Enzyme-related Carboxypeptidase’, Journal of Biological Chemistry, 277: 14838–43.

Vlasak, R, W Luytjes, W Spaan, and P Palese. 1988. ‘Human and bovine coronaviruses recognize sialic acid-containing receptors similar to those of influenza C viruses’, Proceedings of the National Academy of Sciences, 85: 4526–29.

Wrapp, Daniel, Nianshuang Wang, Kizzmekia S. Corbett, Jory A. Goldsmith, Ching-Lin Hsieh, Olubukola Abiona, Barney S. Graham, and Jason S. McLellan. 2020. ‘Cryo-EM structure of the 2019-nCoV spike in the prefusion conformation’, Science, 367: 1260–63.

Wu, Kailang, Guiqing Peng, Matthew Wilken, Robert J. Geraghty, and Fang Li. 2012. ‘Mechanisms of Host Receptor Adaptation by Severe Acute Respiratory Syndrome Coronavirus*’, Journal of Biological Chemistry, 287: 8904–11.

Yan, Renhong, Yuanyuan Zhang, Yaning Li, Lu Xia, Yingying Guo, and Qiang Zhou. 2020. ‘Structural basis for the recognition of SARS-CoV-2 by full-length human ACE2’, Science, 367: 1444–48.

Zheng, Jian, Lok-Yin Roy Wong, Kun Li, Abhishek Kumar Verma, Miguel E. Ortiz, Christine Wohlford-Lenane, Mariah R. Leidinger, C. Michael Knudson, David K. Meyerholz, Paul B. McCray, and Stanley Perlman. 2021. ‘COVID-19 treatments and pathogenesis including anosmia in K18-hACE2 mice’, Nature, 589: 603–07.

Zhu, Na, Dingyu Zhang, Wenling Wang, Xingwang Li, Bo Yang, Jingdong Song, Xiang Zhao, Baoying Huang, Weifeng Shi, Roujian Lu, Peihua Niu, Faxian Zhan, Xuejun Ma, Dayan Wang, Wenbo Xu, Guizhen Wu, George F. Gao, and Wenjie Tan. 2020. ‘A Novel Coronavirus from Patients with Pneumonia in China, 2019’, New England Journal of Medicine, 382: 727–33.

Zoufaly, Alexander, Marko Poglitsch, Judith H. Aberle, Wolfgang Hoepler, Tamara Seitz, Marianna Traugott, Alexander Grieb, Erich Pawelka, Hermann Laferl, Christoph Wenisch, Stephanie Neuhold, Doris Haider, Karin Stiasny, Andreas Bergthaler, Elisabeth Puchhammer-Stoeckl, Ali Mirazimi, Nuria Montserrat, Haibo Zhang, Arthur S. Slutsky, and Josef M. Penninger. 2020. ‘Human recombinant soluble ACE2 in severe COVID-19’, The Lancet Respiratory Medicine, 8: 1154–58.

